# The Anaphase Promoting Complex/ cyclosome co-activator, Cdh1, is a novel target of human Papillomavirus 16 E7 oncoprotein in cervical oncogenesis

**DOI:** 10.1101/2021.11.05.463553

**Authors:** Neha Jaiswal, Deeptashree Nandi, Pradeep Singh Cheema, Alo Nag

## Abstract

The transforming properties of the high risk human papillomavirus E7 oncoprotein are indispensable for driving the virus life cycle and pathogenesis. Besides inactivation of retinoblastoma (Rb) family of tumor suppressors as part of its oncogenic endeavors, E7-mediated perturbations of eminent cell cycle regulators, checkpoint proteins and proto-oncogenes are considered to be the tricks of its transformative traits. However, many such critical interactions are still unknown. In the present study, we have identified the anaphase promoting complex/ cyclosome (APC/C) co-activator, Cdh1, as a novel interacting partner and a degradation target of E7. We found that HPV16 E7-induced inactivation of Cdh1 promoted abnormal accumulation of multiple Cdh1 substrates. Such a mode of deregulation possibly contributes to HPV-mediated cervical oncogenesis. Our mapping studies recognized the carboxyl-terminal zinc finger motif of E7 to associate with Cdh1 and interfere with the timely degradation of FoxM1, a *bona fide* Cdh1 substrate and a potent oncogene. Importantly, the E7 mutant with impaired interaction with Cdh1 exhibited defects in its ability for overriding typical cell cycle transition and oncogenic transformation, thereby validating the functional and pathological significance of the E7-Cdh1 axis during cervical carcinoma progression. Altogether, the findings from our study discover a unique nexus between E7 and APC/C-Cdh1, thereby adding to our understanding of the mechanism of E7-induced carcinogenesis and provide a promising target for the management of cervical carcinoma.

## INTRODUCTION

Cervical carcinoma ranks fourth both in terms of incidence and mortality globally with an estimated 570000 cases, 311000 deaths and less than a 5% survival rate (1). More than 99% of cervical cancer cases are attributed to infection caused by the human papillomavirus (HPV), which has been recognized as the major etiological factor associated with cervical carcinogenesis (2, 3). HPV is a small, double-stranded DNA virus that recruits and modulates the host machinery and functionally abrogates cellular inhibitory factors for viral replication and propagation (4, 5). The HPV encoded viral oncoproteins E6 and E7 have tremendous implications in the induction and maintenance of the transformed phenotype of cervical cancer cells (6). HPV16 E7 is a multifunctional protein, known to bind to and modulate the tumor suppressor factor Rb through the host ubiquitin proteasomal system (UPS) (7–9). E7 controls innumerable oncogenic activities like deregulation of apoptosis, inhibition of cell cycle arrest in response to differentiation or DNA damage, initiation of genomic instability through induction of centrosome duplication errors as well as other mechanisms to establish an environment conducive for viral survival and pathogenesis (10).

The cellular UPS is an elaborate system for organized protein degradation that is critical for maintaining homeostasis among numerous biological functions including accurate cell cycle transition (11). Extensive studies have suggested the key role of cell cycle governed E3 ligases, such as S-phase kinase-associated protein 2 (Skp2) and Cdh1, in the degradation of negative modulators of cell cycle like cyclin/cyclin-dependent kinase (Cdk) inhibitors, p27 and p21, with the help of the UPS (12). Owing to its vital involvement, many viruses are known to hijack this machinery to their advantage (13). The anaphase-promoting complex/cyclosome (APC/C), a canonical E3 ubiquitin ligase complex, targets a spectrum of cell cycle-related proteins, such as cyclins, Polo-like kinase 1 (Plk1) and Skp2, for ubiquitination and subsequent proteasomal degradation in late mitosis and during G1 phase (14). Temporal cell cycle specificity of the APC/C activity is directed mainly through the synchronized binding of two co-activators, Cdc20 and Cdh1 (15, 16). Both Cdc20 and Cdh1 contain WD40 repeats in their C-termini that mediate protein-protein interactions. APC/C has the ability to target an impressive number of substrates involved in a variety of functions central to the cell cycle, like mitotic kinases Aurora kinase A and B, DNA replication, such as Chromatin licensing and DNA replication factor 1 (Cdt1) and Geminin, nucleotide biosynthesis including thymidine kinase and ribonucleotide reductase, and many more (12). While APC/C-Cdc20 functions between metaphase and anaphase, APC/C-Cdh1 mainly exerts its activity from anaphase to cytokinesis until G1 phase. Thus, the explicit management of APC/C-Cdh1 activity is necessary to safeguard normal cell cycle progression and genomic integrity. Interestingly, atypical functioning of Cdh1 is witnessed in a number of cellular disorders including cell cycle defects, impaired cytokinesis and DNA re-replication, leading to overall genomic instability (17).

Not surprisingly, viral factors often exploit the APC/C and disrupt its function with regard to its ubiquitinating ability (18–20). This underscores the potential of APC in driving viral replication and pathogenesis. A striking example is inhibition of APC/C by the Human Cytomegalovirus (HCMV), leading to its dissociation from the positive G_0_/ G_1_ regulator Cdh1 and resulting in accumulation of a plethora APC substrates (21, 22). Involvement of E3 ubiquitin ligases in HPV-induced carcinogenesis has also become increasingly apparent although the mechanism remains unclear. A very fascinating paradigm is that of the high risk HPV E2 protein, which interacts with both the APC activators, Cdh1 and Cdc20, culminating in the buildup of APC substrates and redistribution of Cdh1 into insoluble cytoplasmic aggregates (23). This may explain the fundamental trigger for genomic instability linked with this virus. Of note, E7 expressing cells exhibited elevated levels of early mitotic inhibitor 1 (Emi1), a well-known negative modulator of APC/C-Cdh1 activity (24). Most importantly, HPV16 E6 and E7 act as viral substrate adaptors for the cellular ubiquitin ligases E6AP and Cullin2 to induce proteasome-mediated degradation of p53 (25) and Rb (26) tumor suppressors, respectively. Moreover, we and others have shown the accumulation of the recognized Cdh1 substrate, FoxM1, in HPV positive cervical cancer cells (27). These findings prompted us to examine the possibility if E7 physically interacts with Cdh1 to achieve the hyper-proliferative phenotype in cervical cancer cells. In the present work, we tested the hypothesis that E7 facilitates Cdh1 inactivation, thereby favoring cancerous cell proliferation and metastasis. Our study showed that E7’s interaction with Cdh1 tags it for inappropriate degradation *via* the UPS. Consequently, E7 disturbed the regulated expression of a vast number of Cdh1 substrates. By using Cdh1-interaction defective E7 mutants, we showed that the incapability of E7 to deregulate Cdh1 diminished the oncogenic characters of cervical cancer cells. In a nutshell, our findings underscore a possible mechanism for elevated levels of FoxM1 in HPV infected cervical cancer cells and illustrates that E7 has distinctive mechanism to inactivate the APC/C through proteasomal degradation of Cdh1.

## RESULTS

### HPV16 E7 interacts with Cdh1 *in vivo*

Previous studies have identified multiple viral proteins that tamper with the functions of APC components for their own replication benefits (28–30), implicating the potential role of the latter in virus-induced tumorigenesis. To illuminate a possible connection between E7 oncoprotein and APC co-activator Cdh1, HEK293T cells were transiently transfected with plasmid expressing Myc-E7 either in combination with empty vector or Flag-Cdh1 expression constructs while Flag-p53 was used as an additional negative control. Co-immunoprecipitation (Co-IP) using anti-Flag antibody showed co-immunoprecipitation of Flag-Cdh1 and Myc-E7 (Fig. 1A). This unveiled a physical interaction between E7 and Cdh1. This interaction was further validated at the endogenous level using extracts of CaSki cells, a cervical carcinoma cell line that contains integrated HPV16 DNA (Fig. 1B). Reciprocal IP using anti-E7 antibody further corroborated this interaction (Fig. 1C). Subsequently, co-expression of DsRED-E7 and EGFP-Cdh1 recombinant constructs in HEK293T cells displayed a clear overlap in the localization of Cdh1 and E7 (68.3% colocalization) in the nuclear compartment (Fig. 1D, E, Supplementary Table 1, Supplementary Fig. 1) with a significant rate of co-localization. Altogether, E7 was established to physically associate with cellular Cdh1.

**Fig. 1.**
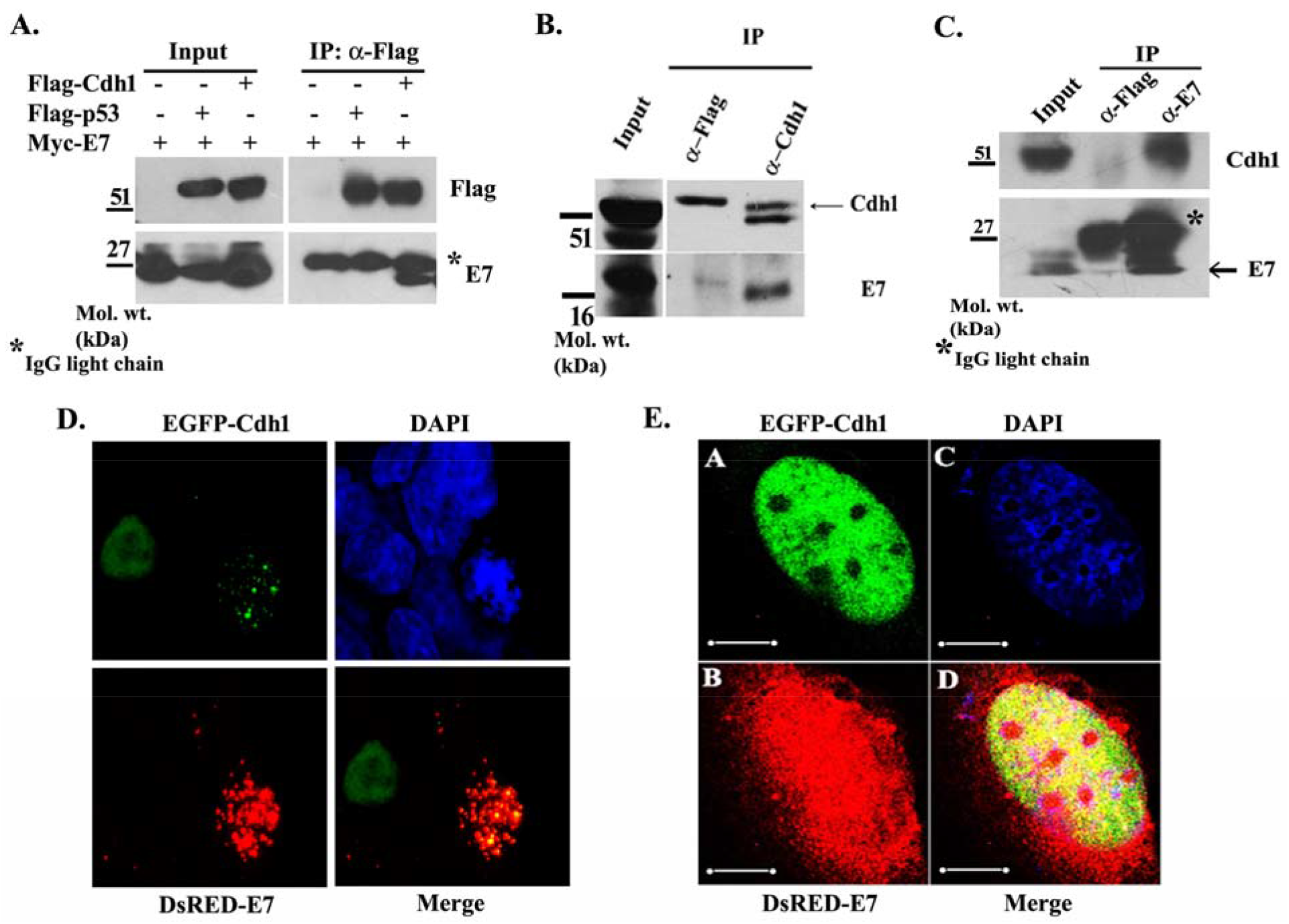
HPV16 E7 associates with Cdh1. (A) E7 interacts with Cdh1 *in vivo*. HEK293T cells were transfected with Myc-E7 in combination with either empty vector, Flag-p53 (negative control bait) or Flag-Cdh1. Cells were lysed and equal amounts of proteins were subjected to co-IP with anti-Flag antibody followed by SDS-PAGE and immunoblotting with anti-E7 and anti-Flag antibody. (B) CaSki cell lysates were immunoprecipitated with control (anti-Flag) or anti-Cdh1 antibody, followed by immunoblotting with E7. (C) Reciprocal IP in CaSki cell lysates was performed with control (anti-Flag) or anti-E7 antibody, followed by immunoblotting for endogenous Cdh1. (D-E) Cdh1 co-localizes with E7. HEK293T cells were co-transfected with EGFP-Cdh1 and DsRED-E7. Post-transfection, cells were fixed and counterstained with DAPI and images were captured using confocal microscope (D, 40x and E, 100x). Scale bar, 10 *μ*m.

### Cdh1 is inactivated in E7-expressing cells

In view of the function of E7 in degradation of host tumor suppressor proteins (9, 31, 32), we wanted to investigate the outcome of E7 on Cdh1 stability. Hence, we examined the level of endogenous Cdh1 protein in C33A cells stably expressing HPV16 E7. The presence of E7 in these cells was verified through detection of E7 mRNA using RT-PCR (Fig. 2A). Over-expression of E7 culminated in a clear reduction of Cdh1 level with a concomitant increase in the level of FoxM1, which is known to be negatively regulated by Cdh1 (Fig. 2B). Conversely, silencing of E7 in CaSki cells resulted in the accumulation of cellular Cdh1 (Fig. 2D). The results revealed an inverse correlation between E7 and Cdh1 and suggested a pivotal role of E7 in altering steady state of Cdh1 protein. We next wished to address the functional significance of the interaction between Cdh1 and E7. As part of cell cycle control, Cdh1, in association with APC, oversees the turnover of a number of substrates (33). We found that degradation of important mitotic substrates of Cdh1 like Plk1, Aurora A and Skp2 was compromised in the presence of E7 (Fig. 2C). Expression of E7 also facilitated elevated levels of other Cdh1 substrates, including the DNA replication factor Cdt1 and Geminin, whose degradation releases Cdt1 from inhibition during late mitosis (Fig. 2C). These results were in accordance with previous reports (24, 34–37) and further denoted the important role of E7 in manipulating Cdh1 level in cancer cells. In agreement, E7 depletion in CaSki cells led to a notable decrease in the levels of Cdt1, Geminin, FoxM1, Plk1, Skp2, CyclinB1 and Aurora B Kinase, thus reflecting the restoration of their timely regulation by Cdh1 (Fig. 2D). The findings designate E7-mediated inactivation of Cdh1 as a novel mechanism for HPV pathogenesis.

**Fig. 2.**
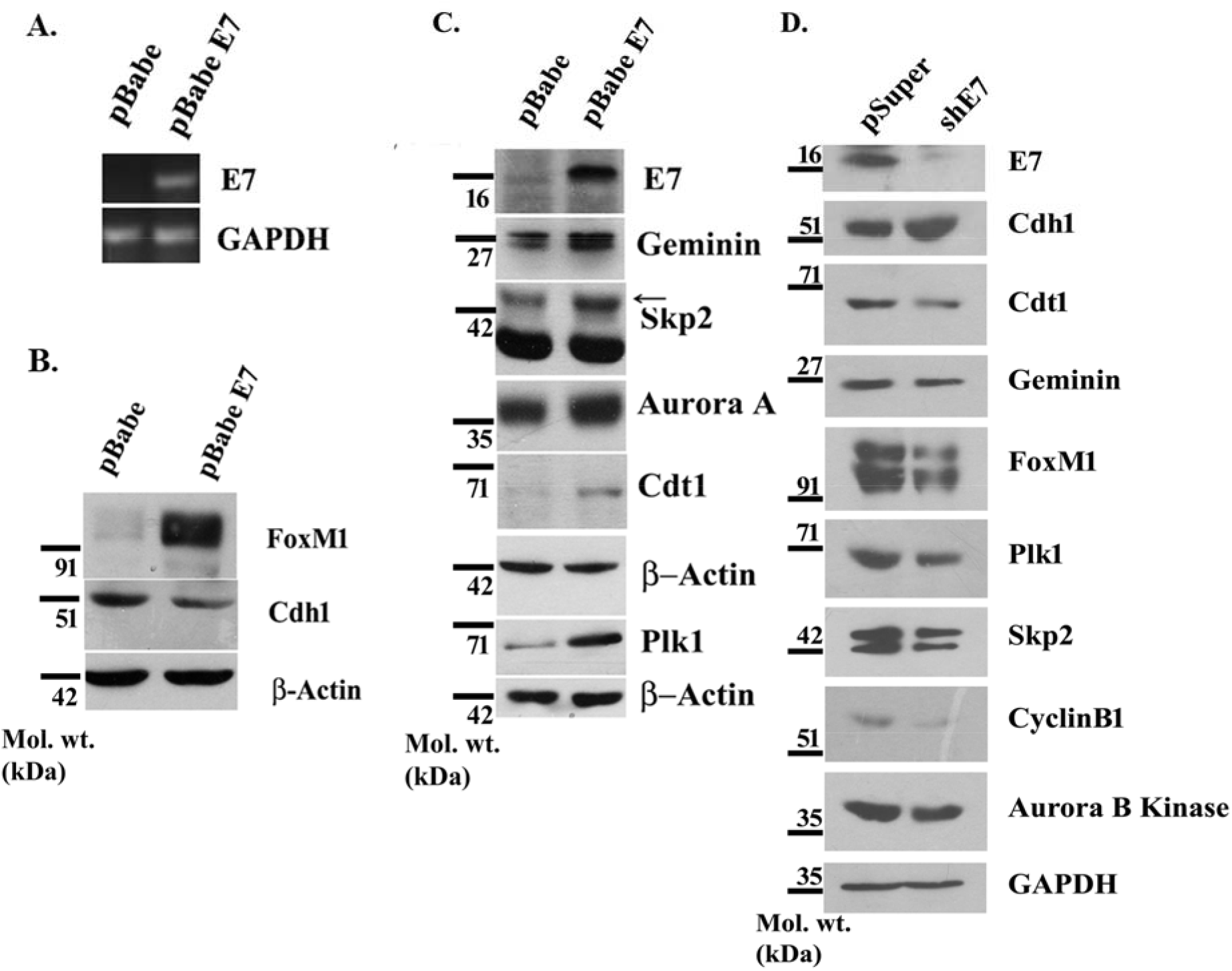
HPV16 E7 alters steady state levels of Cdh1. (A) Generation of C33A cells stably expressing E7. Total RNA was extracted from C33A cells stably expressing HPV16 E7. E7 and GAPDH mRNA levels were analyzed by RT-PCR. (B) Cdh1 is diminished by E7. Cell lysates from C33A cells with stable expression of pBabe control vector or pBabe E7 cells were subjected to Western blotting with antibodies against Cdh1, FoxM1 and β-Actin. (C) HPV16 E7 interferes with the down-regulation of multiple Cdh1 substrates. Protein extracts were prepared from C33A cells stably expressing pBabe control vector or pBabe E7 and steady state levels of E7, Geminin, Skp2, Aurora A, Cdt1 and Plk1 were analyzed by Western blotting with specific antibodies. β-Actin was used as a loading control. (D) Depletion of E7 stabilized Cdh1. Western blot analysis of CaSki cells expressing either control vector pSuper retro puro or shE7 with antibodies specific for E7, Cdh1, Cdt1, Geminin, FoxM1, Plk1, CyclinB1, Skp2 and Aurora B Kinase. GAPDH served as a loading control.

### HPV16 E7 promotes degradation of Cdh1 *via* the ubiquitin proteasome pathway

To further explore the influence of E7 on Cdh1 protein stability, HEK293T cells were transfected with Myc-Cdh1 alone or in combination with Myc-E7 followed by cycloheximide (CHX) chase assay. We observed a sharp decline the half-life of Cdh1 from 3.5 h to less than 1 h in the presence of Myc-E7 (Fig. 3A, B). Similar observations in C33A cells stably expressing HPV16 E7 (Fig. 3C, D) strongly substantiated our hypothesis that E7 destabilizes Cdh1. Given the well-documented role of E7 in the degradation of Rb (9), the diminished expression of Rb in the C33A stable cell line confirmed the presence of E7. Past research has implicated the role of E7 in manipulating the UPS for aberrant degradation of host proteins (26). Therefore, we asked whether proteasomal block could stabilize Cdh1 in presence of E7. Treatment of E7 over-expressing HEK293T cells with MG132 resulted in enhanced Cdh1 protein level (Fig. 3E), endorsing that E7-related suppression in Cdh1 expression was, at least partially, due to proteasomal degradation. The disappearance of E7 in Fig. 3E (second lane) reflects the short half-life of E7, which was restored upon treatment with MG132 (Fig. 3E, third lane). This implied the involvement of UPS in the stability of E7, which was in concordance with previous reports (38). Finally, our *in vivo* ubiquitination assay demonstrated a significant increase in the level of ubiquitin-conjugated Cdh1, which appeared as a smear of high molecular weight bands, in presence of E7 (Fig. 3F). Such pattern was lacking in control cells. Together, these results authenticated that E7 stimulates ubiquitin tagging of Cdh1, thus priming it for the proteasomal degradation.

**Fig. 3.**
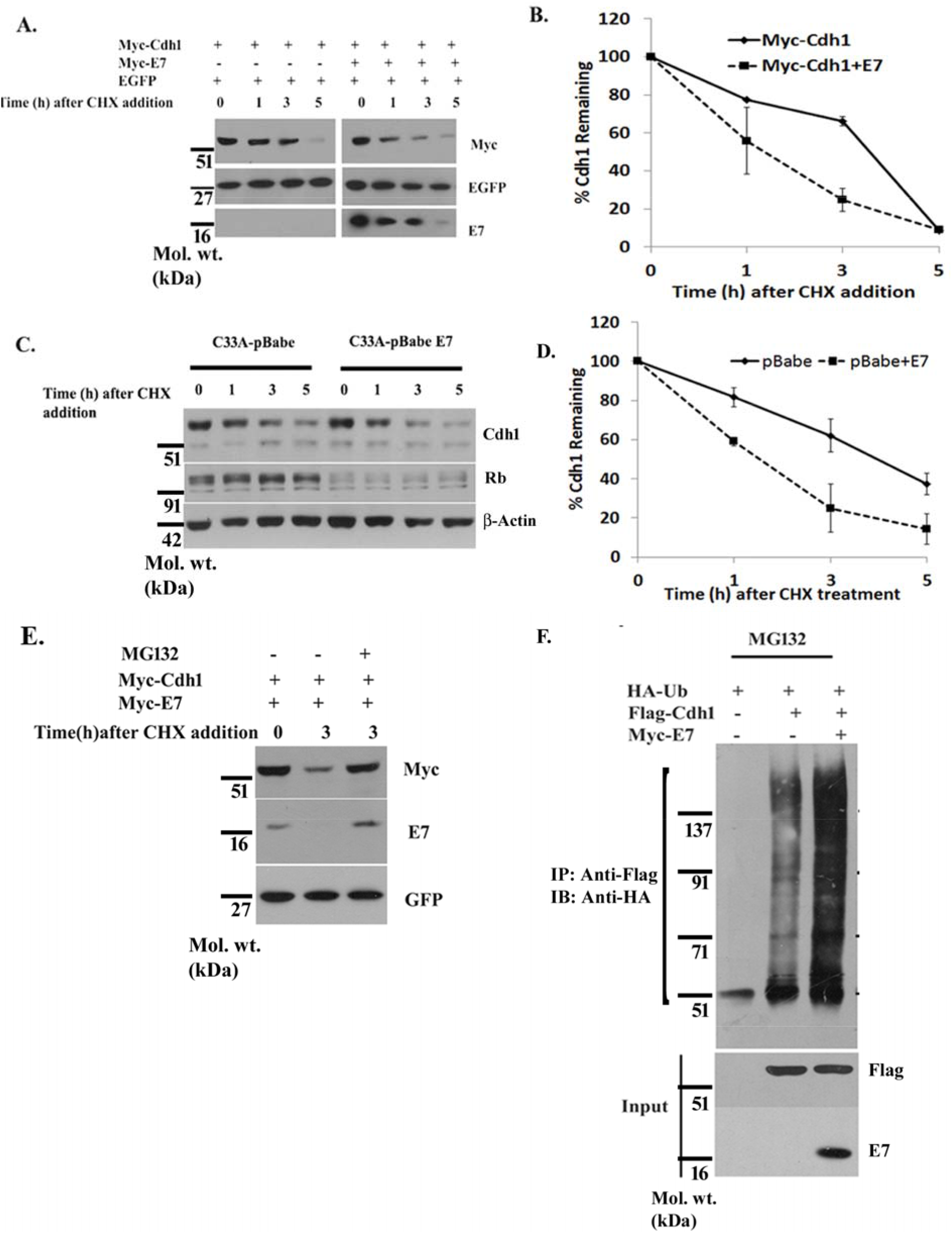
HPV16 E7 oncoprotein destabilizes Cdh1. (A) HEK293T cells were transfected with Myc-Cdh1 alone or with Myc-E7 expression plasmids. Forty hours after transfection, cells were incubated with CHX (100 μg/ml) for indicated time-points and cell lysates were immunodetected with anti-Myc and anti-E7 antibodies. GFP was used as a transfection control (B) Graphical representation of the intensities of Cdh1 bands quantified using densitometric analysis as mentioned in experimental procedures. (C) C33A cells stably expressing HPV16 E7 were subjected to CHX half-life assay at indicated time-points. Western blot analysis was performed with antibodies against endogenous Cdh1, Rb and β-Actin. The presence of E7 in the stable line was verified by monitoring Rb expression. (D) The intensities of Cdh1 bands were quantified using densitometric analysis as mentioned in experimental procedures and graphically depicted. (E) HPV16 E7 stabilizes Cdh1 in the presence of MG132. HEK293T cells transfected with Myc-Cdh1 and Myc-E7 were subjected to MG132 treatment for 5 h. CHX was added 2 h post-addition of MG132 for another 3 h. Equal amount of cell lysates were subjected to SDS-PAGE followed by western blotting with Myc, E7 and GFP antibodies. (F) HPV16 E7 enhances the ubiquitination of Cdh1 *in vivo.* HEK293T cells were transfected with the expression plasmids HA-Ub, Flag-Cdh1 and Myc-E7 as indicated. Transfected cells were treated with MG132 for 5 h. Total cell extracts were then subjected to IP using anti-Flag antibodies under denaturing conditions. Poly-ubiquitinated Cdh1 was detected using anti-HA antibody. The expression of Flag-Cdh1 and Myc-E7 was detected in whole cell lysate (input).

### Cdh1 medial domain is important for E7-mediated destabilization

Earlier reports indicate that Cdh1 contains an N-terminal C-box (39), critical for interaction of the APC/C and its E3 ligase activity (40, 41), and a more prominent C-terminal domain with WD40 repeats (42, 43) provides a substrate-binding platform. In order to gain insight into the domain responsible for E7-mediated destabilization of Cdh1, we engineered a set of Cdh1 deletion mutants, including N-terminal Cdh1 (1-124aa) domain, C-terminal domain (201-493aa) and a middle domain (1-200aa) (Fig. 4A). Using CHX chase assay, we assessed the stability of these Cdh1 deletion mutants in cells ectopically expressing E7. As shown in Fig. 4B and C, the Cdh1 mutant encompassing the middle domain flaunted a significant drop in its half-life while the remaining mutants were relatively stable. Next, the Cdh1 mutants that conferred resistance to E7-mediated degradation were further checked for their ubiquitination abilities in the presence of E7. As expected, wild type Cdh1 displayed a normal ubiquitination pattern as seen before (Fig. 3F) but Cdh1 deletion mutants (1-124aa and 201-493aa) presented a defect in their ubiquitination pattern (Fig. 4D). Taken together, our mutational studies revealed that the region between residues 124 and 200aa of Cdh1 protein is sensitive and imperative to E7-induced Cdh1 interaction and degradation.

**Fig. 4.**
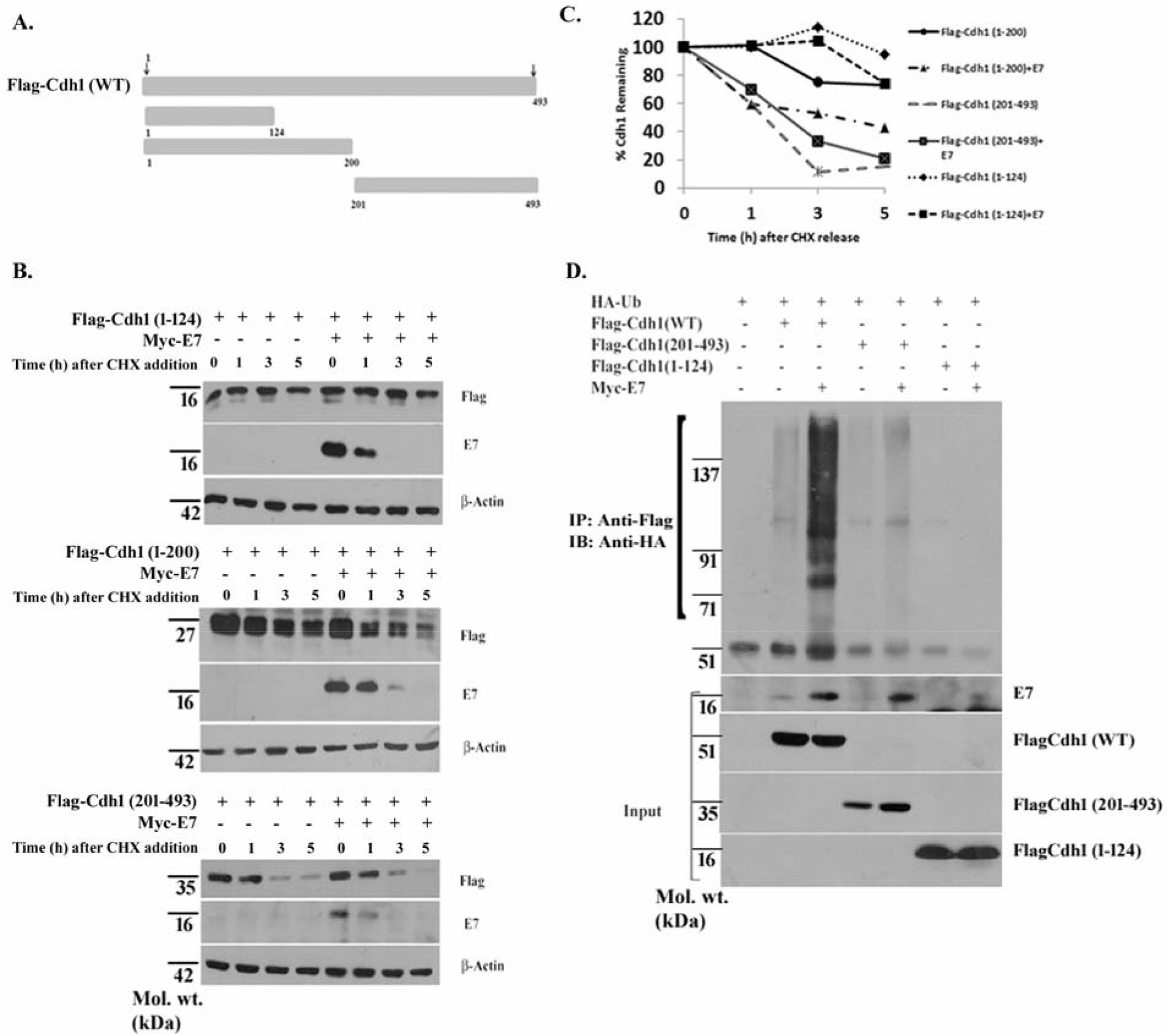
Medial region of Cdh1 is required for E7-induced inactivation. (A) Schematics of Cdh1 deletion mutants. (B) Analysis of protein stability for Cdh1 mutants in the presence of E7 using CHX chase assay. Decay rate of Cdh1 protein was followed till indicated time-points. (C) The intensities of Cdh1 bands were quantified and normalized using densitometric analysis as mentioned in experimental procedures and graphically represented. (D) Defect in the ubiquitination pattern of Cdh1 mutants. HEK293T cells were transfected with either HA-Ub alone or in combination with wild type Cdh1 or its deletion mutants (1-124aa) and (201-493aa) in the absence and presence of Myc-E7. Transfected cells were treated with MG132 for 5 h. Total cell extracts were then subjected to IP using anti-Flag antibody under denaturing conditions. Poly-ubiquitinated Cdh1 was detected using anti-HA antibody.

### E7 carboxyl-terminal domain destabilizes Cdh1

Delineation of the E7 domain important for its association with Cdh1 was achieved by employing three deletion mutants of DsRED-E7 containing the N-terminal E7 (1-15aa) conserved region (CR) 1 homology domain, the medial E7 (16-37aa) CR2 homology domain that possesses the Rb-binding LXCXE motif and the C-terminal (38-98aa) cysteine rich zinc-finger motif (44, 45) (Fig. 5A). Analysis of Cdh1 protein turnover ascertained that the C-terminal E7 mutant dramatically diminished Cdh1 stability (Fig. 5B, C). In contrast, Cdh1 half-life remained unaltered in the presence of the other two E7 mutants (Fig. 5B, C). In agreement with the stability study, our co-IP result identified the C-terminal region of E7, encompassing 38-98aa residues, as part of the essential interacting interface with Cdh1 (Fig. 5D). Next, we sought the ability of these mutants to impart proteolytic degradation of Cdh1. Consistent with our findings from interaction and half-life studies, the C-terminus of E7 promoted strong ubiquitination of Cdh1 while the other mutants failed to do so (Fig. 5E). Our localization study detected all three E7 mutants to cohabit with Cdh1 (Fig. 5F, Supplementary Table 1, Supplementary Fig. 1). This insinuated that simply co-existing together does not allow E7-mediated repression of Cdh1. With the goal to further pinpoint the Cdh1-interaction motif of E7, we employed the E7 C58Gly mutant, where the cysteine residue within the zinc-binding motif in the E7 CR3 domain was mutated. While a strong interaction was visible between Flag-Cdh1 and Myc-E7 following IP with anti-Myc antibody, interestingly, the C58Gly E7 mutant showed little or no binding with Cdh1 (Fig. 5G). This implied the significance of the cysteine residue within the zing finger domain of E7 for its steady interaction with Cdh1. Of importance, Cdh1 failed to interact with the p24Gly E7 mutant, which is well-known for its defect in Rb binding (46), hinting at a plausible role of Rb in Cdh1-E7 interaction (Fig. 5G). Therefore, our mutational analysis advocated that the functional abrogation of Cdh1 necessitates an intact carboxyl terminus of E7.

**Fig. 5.**
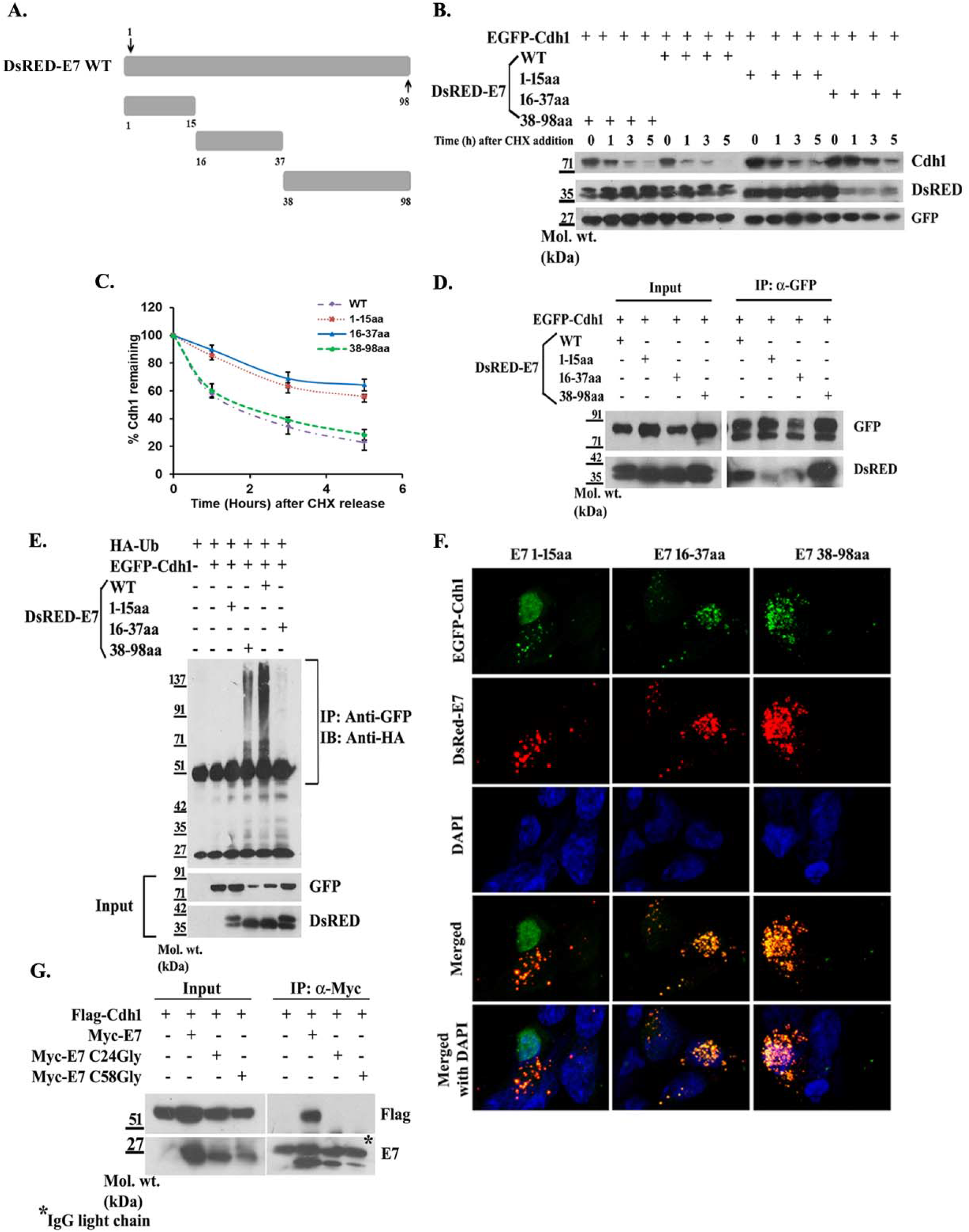
E7 carboxyl terminus is the functional domain mediating Cdh1 degradation. (A) Schematic representation of the E7 deletion mutants. (B) E7 C-terminus induces rapid protein turnover of Cdh1. CHX chase assay of Cdh1 was performed in presence of DsRED-E7 WT and the mutants. Decay rate of Cdh1 protein was followed till indicated time-points. (C) The intensities of Cdh1 bands were quantified and normalized using densitometric analysis as mentioned in experimental procedures and graphically depicted. (D) E7 C-terminus mimics the wild type in its binding with Cdh1. HEK293T cells were over-expressed with EGFP-Cdh1 in combination with either DsRED-E7 WT or its deletion mutants (1-15aa), (16-37aa) or (38-98aa), followed by co-IP using anti-GFP antibody. The mutants containing (1-15aa) and (16-37aa) were found to interact very weakly with Cdh1 whereas the E7 mutant comprising of amino acids 38-98 interacted with Cdh1 to the same extent as the WT. (E) E7 38-98aa mutant prominently ubiquitinates Cdh1. HEK293T cells were transfected with either HA-Ub alone or in combination with DsRED-E7 WT or its deletion mutants in the presence of EGFP-Cdh1. Cells were treated with MG132 for 5 h followed by IP of the whole cell lysates using anti-GFP antibody under denaturing conditions. Poly-ubiquitinated forms of Cdh1 were detected using anti-HA. (F) Co-localization pattern of Cdh1 with the E7 mutants. HEK293T cells were co-transfected with EGFP-Cdh1 and DsRED-E7 (1-15aa), (16-37aa) or (38-98aa). Cells were fixed and nuclei counter-stained with DAPI followed by confocal microscopy. (G) E7 zinc-binding motif interacts with Cdh1. HEK293T cells were transfected with Flag-Cdh1 either alone or in combination with Myc-E7, Myc-E7 p24Gly or Myc-E7 C58Gly expressing plasmids. Following IP with anti-Myc antibody, immunodetection was carried out using anti-Flag antibody.

#### Overexpression of E7 protects oncogenic FoxM1 from Cdh1-mediated regulation

One of the principle substrates of the APC/C-Cdh1 is the oncogenic transcription factor FoxM1 (47, 48), whose physiological level is maintained through its regulated proteolysis in a cell cycle dependent manner by Cdh1 (49). Our previous study has noted high levels of FoxM1 in HPV positive cervical cancer cell lines (27). Having identified Cdh1 as a major target of E7, we sought to examine whether tampering with Cdh1 levels by E7 would have a corresponding outcome on FoxM1 stabilization. Interestingly, introduction of E7 in C33A cells led to a noticeable rarefication in Cdh1 level and an appreciable elevation of FoxM1 (Fig. 6A). Furthermore, intensification in the level of a number of genes downstream of FoxM1, including Plk1, CyclinB1, Skp2, Aurora B Kinase and Centromeric protein A (CENP-A), strengthened our proposed role of E7 in shielding FoxM1 from Cdh1-mediated destabilization (Fig. 6A). Ectopic expression of FoxM1 in combination with control vector or Cdh1 and increasing doses of E7 additionally disclosed partial restoration in FoxM1 level (Fig. 6B). We further tested the possible attenuation of Cdh1 function by E7 through dissociation of the FoxM1-Cdh1 complex through co-IP. Surprisingly, E7 did not disrupt the interaction between FoxM1 and Cdh1 (Fig. 6C), instead a band corresponding to E7 in IP lane was seen. This conveyed that the three proteins may exist together as a trimeric complex and E7 might disable Cdh1 by creating a functionally inactive complex. Our next goal was to investigate the outcome of the E7 mutants on the expression of FoxM1. Over-expression of distinct E7 mutants in HEK293T cells indicated that E7 C-terminus successfully rescued FoxM1 from Cdh1-mediated suppression in a manner similar to full length E7. However, no such restoration was seen in presence of the Cdh1-interaction defective E7 mutants (Fig. 6D, E). This suggested that E7’s interaction with Cdh1 is essential for the viral oncoprotein to deregulate the levels of Cdh1 substrates such as FoxM1, which may be an important mechanism underlying E7-induced transformations.

**Fig. 6.**
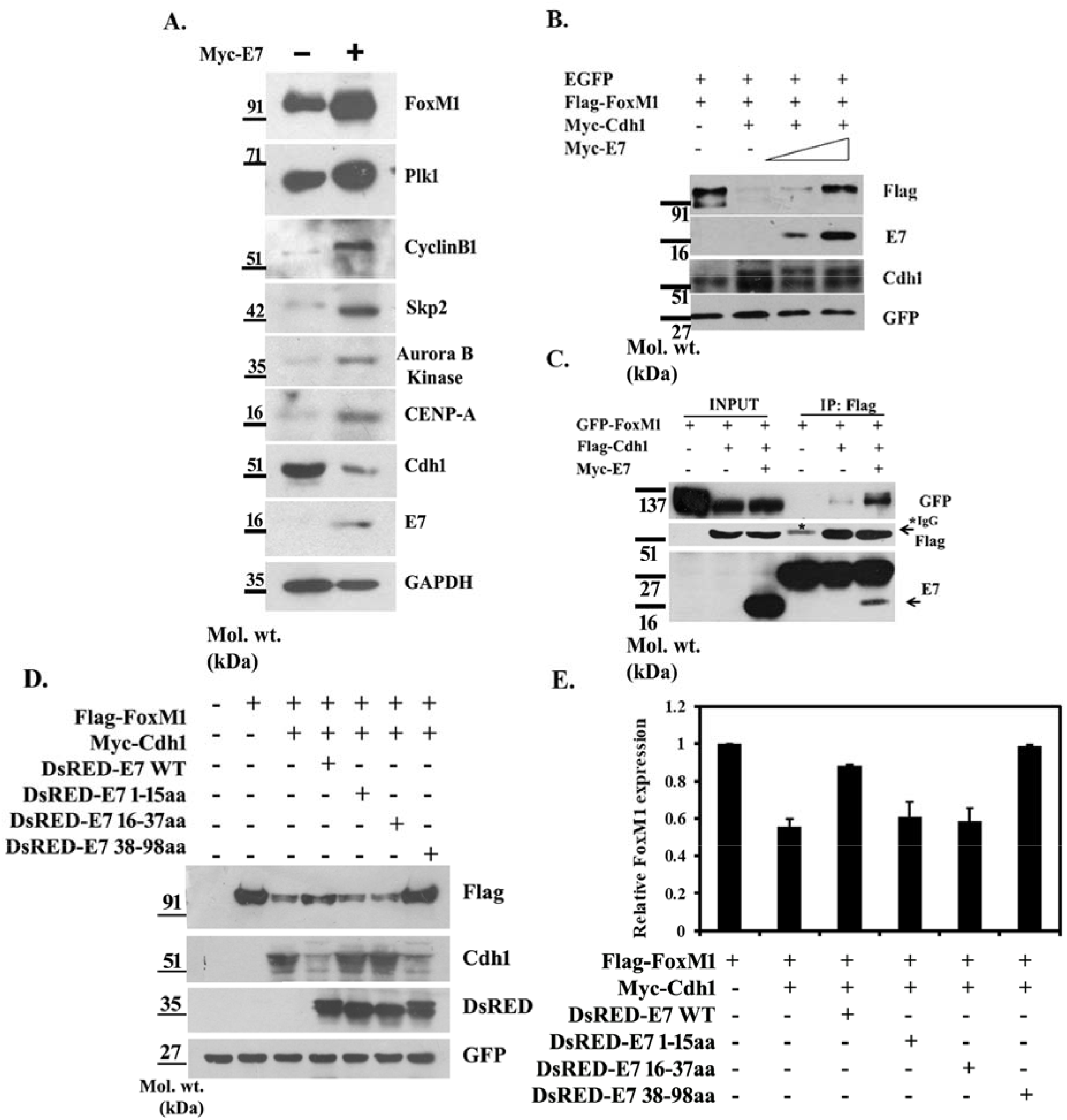
HPV16 E7 interferes with Cdh1-mediated degradation of FoxM1. (A) HPV16 E7 rescues FoxM1 from Cdh1. C33A cells were transiently transfected with either empty vector or Myc-E7 and equal amounts of whole cell lysates were subjected to SDS-PAGE and immunoblot analysis. Expression of E7 was confirmed through detection using antibody specific for E7. Western blotting was carried out against endogenous Cdh1 as well as FoxM1 and its principle downstream effector genes like Plk1, CyclinB1, Skp2, Aurora B Kinase and CENP-A. GAPDH was used as a loading control. (B) Restoration of FoxM1 levels in the presence of HPV16E7. HEK293T cells were co-transfected with Flag-FoxM1 in combination with control vector, Myc-Cdh1 and increasing doses of Myc-E7. Following transfection, cell lysates were subjected to SDS-PAGE followed by immunoblotting with antibodies for Flag, E7, Cdh1 and GFP (transfection control). (C) HPV16 E7 does not perturb the interaction between FoxM1 and Cdh1. HEK293T cells were transfected with either EGFP-FoxM1 alone or in combination with Flag-Cdh1 in the absence and presence of Myc-E7. Whole cell lysates were then subjected to co-IP with anti-Flag antibody followed by immunodetection with antibodies against GFP, Flag and E7 antibody. (D) HEK293T cells were transiently transfected with Flag-FoxM1 either in presence of empty vector or Myc-Cdh1 and DsRED-E7 WT and the mutants. Cells were harvested 48 h later and subjected to SDS-PAGE. Immunoblotting with antibodies specific for anti-Flag, anti-Myc and anti-DsRED revealed a restoration in level of Flag-FoxM1 in the presence of WT and C-terminal E7. GFP was used as a transfection control. (E) The band intensities for Flag-FoxM1 were measured using densitometric analysis and the relative FoxM1 expression (normalized to the control) was graphically presented.

### Cdh1 ablation by E7 delayed mitotic exit and impaired neoplastic transformation

The E7 oncoprotein is documented to promote abnormal cell cycle transition by interacting with and neutralizing key cell cycle related proteins, like Rb, thereby activating proteins responsible for anomalous S phase entry (50). Cdh1 is an essential element that maintains accurate transition of cell cycle phases by maintaining optimal turnover of key cell cycle controllers. To illuminate the novel role of Cdh1 in E7-mediated cell cycle alteration, we studied the expression profile of different cell cycle mediators in C33A cells transiently expressing the E7 mutants. Immunoblotting results showed normal levels of CyclinB1 and CyclinE in presence of the Cdh1-interaction defective E7 mutants while introduction of full length or C-terminal E7 resulted in an abnormal accumulation of these cyclins (Fig. 7A). This implied that altered Cdh1 activity due to its interaction with E7 disturbed the timely degradation of CyclinB1 and CyclinE. Complementing this result, we observed relegated protein levels of CyclinB1 and CyclinE in C33A-E7 stable line with Cdh1 knockdown (Fig. 7B). In continuation, BrdU incorporation assay was performed to examine whether S phase progression is also influenced by the Cdh1-E7 axis. C33A cells stably expressing E7 exhibited increased BrdU incorporation rate (28.8%) compared to the control C33A cells (21.4%) (Fig. 7C, D), in agreement with published report (50). Interestingly, Cdh1 depletion in C33A-E7 stable cells led to further escalation in the BrdU positive population (51.1%), depicted in Fig. 7C and D. Such data supported the established notion that ablation of Cdh1 prolongs S phase in cancer cells (51). Ectopic expression of the Cdh1-interaction defective E7 (1-15aa) mutant also gave a similar rise (45%) in the S phase population (Fig. 7C, D). Furthermore, our cell cycle analysis confirmed that while C33A cells, in general, displayed an average of 17.63% of cells in the S phase, this population nearly doubled to 30% following Cdh1 knockdown in the C33A-E7 stable cell line (Fig. 7E, F). Similarly, in cells expressing E7 (1-15aa) mutant, the S phase cells amounted to approximately 26.3%, thereby corroborating our proposition that E7 hijacked Cdh1 activity to favor untimely mitotic progression. However, with loss in its ability to control Cdh1, E7-expressing cells experienced a prolonged S phase and a delayed entry into mitosis. Therefore, our data represented Cdh1 deregulation as an unequivocal requirement during E7-mediated deviations in the host cell cycle machinery.

**Fig. 7.**
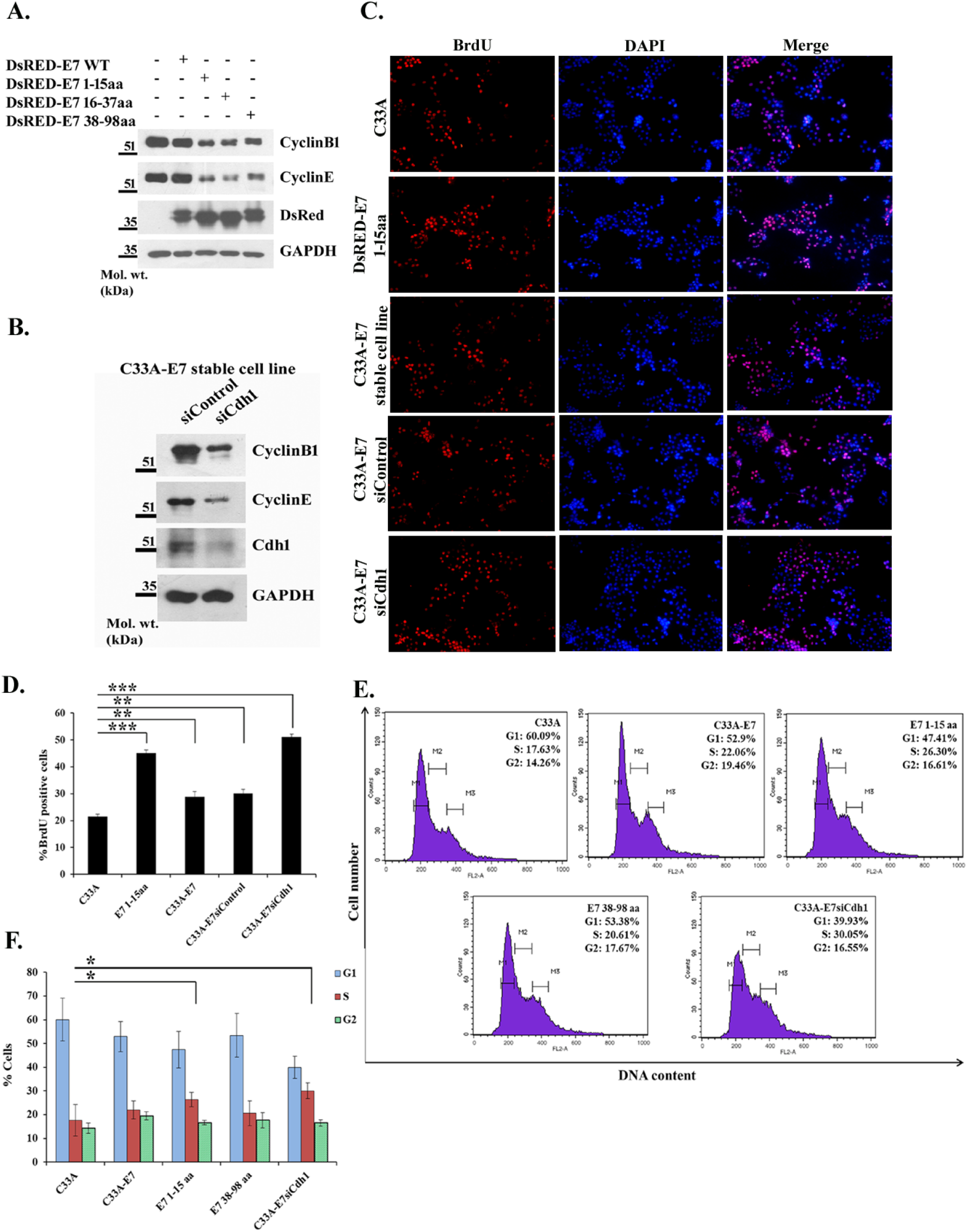
Loss of Cdh1 abrogates E7-related cell cycle deregulation. (A) E7 requires Cdh1 for accumulation of CyclinB1 and CyclinE. C33A cells were transiently transfected with DsRED-E7 WT and its deletion mutants. Equal amount of whole cell lysates were subjected to SDS-PAGE. Western blot analysis was carried out for CyclinB1 and CyclinE. GAPDH was used as a loading control. (B) C33A-E7 line stably expressing the full length E7 was subjected to Cdh1 knockdown using siRNA construct for 48 h followed by SDS-PAGE and immunoblotting with anti-CyclinB1, anti-CyclinE, anti-Cdh1 and anti-GAPDH antibodies. (C) C33A cells were transiently transfected with E7 1-15aa mutant whereas C33A cells stably expressing full length E7 were subjected to Cdh1 knockdown. A scrambled siRNA construct (siControl) was used as a negative control for this purpose. Cells were pulsed with BrdU and incorporation of BrdU was detected through immunofluorescence using an anti-BrdU antibody as has been explained in Materials and methods. The figure presents data representative of results from at least three independent experiments. (D) The number of BrdU positive cells was plotted as percentage of the total number of cells for each set. The means and respective standard deviations are indicated. (E) Flow cytometric analysis of the cell cycle populations. C33A cells transiently expressing E7 1-15aa or 38-98aa mutants and C33A-E7 stable cells silenced of Cdh1 were fixed and stained with propidium iodide as mentioned in Materials and methods and subjected to analysis in the flow cytometer. The figure presents the number of cells in various phases of the cell cycle, detected by the DNA content of each cell. The data are representative of results from at least three independent experiments. (F) The percentage of cells in each of G1, S and G2 phases has been graphically presented for the different sets. The means and respective standard deviations are indicated. Statistical significance for respective graphs was determined by Student’s *t* test. *, *P* < 0.05 and *P* > 0.01; **, *P* < 0.01 and *P* > 0.001; ***, *P* < 0.0001, compared to the control.

HPV16 E7 reportedly engages different host proteins for epithelial-to-mesenchymal transition (EMT), an important phenomenon during cervical tumorigenesis, which is accompanied by increased cell migration and invasiveness (52). To assess whether Cdh1 is recruited by E7 during this process, we transfected C33A cells with either the N- or the C-terminal E7 mutant and transiently silenced Cdh1 in E7-expressing C33A stable cell line. Thereafter, we examined their effects on cell migration and invasion using wound-healing assay and invasion assay, respectively. Cells stably expressing full length E7 healed effectively with less than 30% of the original gap remaining after 72 h (Fig. 8A, B), suggesting that E7 positive cervical carcinoma cells migrated more extensively compared to control cells. The C-terminal mutant of E7 also exhibited significant (45% wound remaining at 72 h) gap closure (Fig. 8A, D), comparable to the migratory index of full length E7. On the contrary, a significant decline in the gap closure was observed upon Cdh1 silencing in C33A-E7 cells (Fig. 8A, F) or expression of the E7 (1-15aa) mutant in C33A cells (Fig. 8A, C). The results implied that E7 needs to interact with and deregulate Cdh1 for enhanced tumor cell proliferation. Similar results were obtained for invasion assay, where cells stably expressing full length E7 displayed a sharp increase (approximately 2-fold) in their invasive potential whereas loss of Cdh1 or presence of the Cdh1-inactivation deficient E7 mutant dramatically decreased the invasiveness (Fig. 8G, H). Collectively, these findings clearly suggest that HPV-related transformation depends on the ability of E7 to commandeer and override normal functions of Cdh1.

**Fig. 8.**
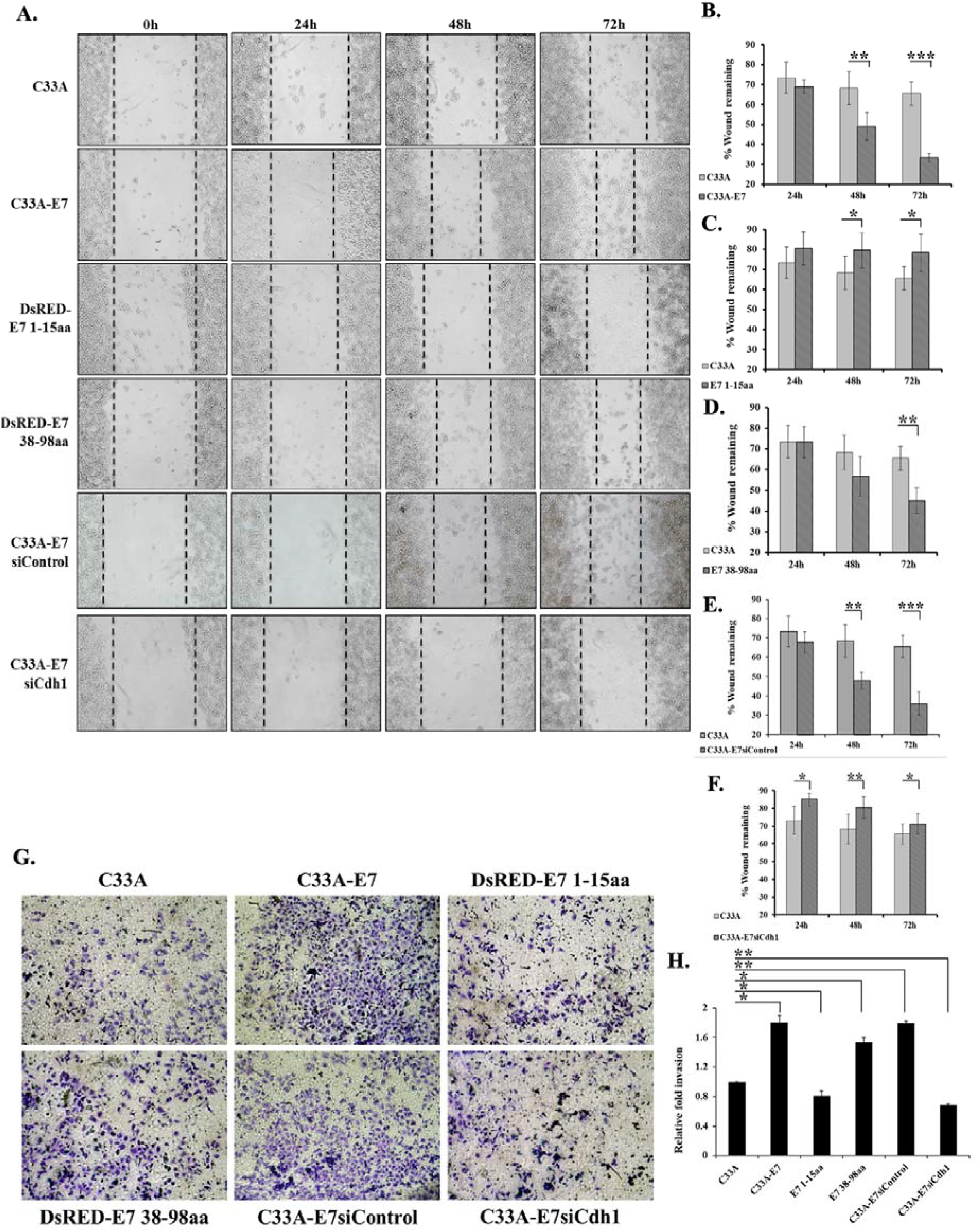
E7-induced neoplastic transformation is perturbed upon Cdh1 depletion. (A) C33A cells were transiently transfected with either E7 1-15aa or 38-98aa mutants while siRNA-based Cdh1 knockdown was done in C33A cells stably expressing WT E7. These cells were thereafter seeded so that a confluent monolayer is attained the following day. Thereafter, a wound was introduced using a 20 μl tip and the gap closure was photographed at 0, 24, 48 and 72 h. The average relative distances between the edges of the wounds at least at six different points are shown. The figure displays data representative of results from at least three independent experiments. (B-F) The percent wound remaining has been measured and plotted for each time point with respect to the original wound at 0 h. The comparison is presented between control C33A cells and cells expressing either WT E7, E7 1-15aa or E7 38-98aa mutants and between C33A cells with C33A-E7 stable line treated with either siControl or siCdh1 construct. The means and respective standard deviations are indicated. (G) The Matrigel invasion assay was performed in C33A cells stably expressing E7 WT or transiently expressing E7 1-15aa or 38-98aa and C33A-E7 stable line with siRNA-based Cdh1 knockdown as described in Materials and Methods. Forty eight hours after transfection, the invaded cells were stained with crystal violet, photographed and counted. The figure presents data representative of results from at least three independent experiments. (H) Bar graph depicting fold change of invading cells relative to the control set, calculated from at least six independent fields of view. The means and respective standard deviations are indicated. Statistical significance for respective graphs was determined by Student’s *t* test. *, *P* < 0.05 and *P* > 0.01; **, *P* < 0.01 and *P* > 0.001; ***, *P* < 0.0001, compared to the control.

## DISCUSSION

The APC/C-Cdh1 complex is a climacteric factor indispensible for maintaining physiological equilibrium within the host system (12, 17, 53). Mutational and knockout studies have recognized the tumor suppressive role of this axis (33, 53–55) and alteration to its functions have been implicated in several human carcinomas (33). Contemporary studies have shown HPV16 E7 to destabilize numerous tumor suppressors, particularly by exploiting the host UPS. E7 protein of assorted HPV types and HPV16 E6 are reported to bind to a wide array of UPS factors, such as RING-type Ub ligases and the DUB Ub-specific protease 15 (56). Of note, E7-mediated Rb degradation involves the cullin 2 ubiquitin ligase complex (26). Perturbation of the DREAM complex resulting in up-regulation of cellular oncogenic factors is yet another example of E7-related host function remodeling during cervical carcinogenesis (57). In sync, our evidences hint at a plausible role of HPV16 E7 in disrupting the activity of Cdh1 to impart the diverse tumorigenic functions in E7-transformed cancer.

In this study, we testify, for the first time, that Cdh1 interacts with HPV16 E7 (Fig. 1A). Our mapping studies showed that the CR3 domain of E7 is sufficient for its interaction with Cdh1 (Fig. 5). In addition, we showed that the medial portion of Cdh1 associates with E7 (Fig. 4). Amino acid analysis of this region revealed multiple lysine residues that could serve as docking sites for ubiquitin attachment, thus explaining the sensitivity of this domain towards E7-mediated proteolysis (58). Of note, we identified C58 as an essential residue for the interaction between Cdh1 and HPV16 E7 as mutation of this residue completely abolished their interaction (Fig. 5G). The E7 CR3 domain serves as a pivotal platform for interaction and perturbation of key cellular factors with implications in the transformative potential of the viral oncoprotein (59–61). Importantly, although the most thoroughly documented Rb-binding pocket of E7 is its high-affinity binding site in the CR2 region, crystallographic and mutational analysis have pointed to a secondary binding site for Rb in the CR3 domain of E7 (62–64). Intriguingly, this interaction was shown to affect Rb degradation and deregulation comparable to wild-type E7. Surprisingly, although a very weak binding was observed between Cdh1 and the E7 residues between 16-37aa, Cdh1 failed to interact with the Rb-defective C24Gly E7 mutant (Fig. 5G). These findings raise the possibility that both Cdh1 and Rb may share a common interaction interface with E7 or E7 bound to Rb provides the necessary conformation that exposes the residues required for E7-Cdh1 interaction. Thorough research initiatives will enable further insight into the plausible role of Rb in E7-Cdh1 functions.

A phenomenal finding was the association between Cdh1 and E7 at the endogenous level in HPV positive cervical cancer cells (Fig. 1B, C). This indicated towards the relevance of this interaction in cervical tumors of HPV positive patients, thus giving prominence to the Cdh1-E7 complex in E7-linked oncogenic transformation. Clinical assessment of the correlation of this association with cervical cancer prognosis is likely to benefit HPV positive patients as targeting this axis may improve overall survival. Furthermore, we found preferential co-localization between E7 and Cdh1 in the nuclear compartment (Fig. 1D, E). A previous report proposed that nuclear localization of E7 is pivotal for its pathological functions (65). Our finding of co-habitation of Cdh1 and E7 may, thus, serve as an important mechanism in HPV propagation and oncogenesis. Importantly, our results hinted towards an inverse correlation between E7 and Cdh1 (Fig. 2). Delineation of the molecular mechanism unveiled that E7 augmented the proteolytic turnover and proteasomal degradation of Cdh1 (Fig. 3). Indeed, Cdh1 down-regulation has been detected in human cancer of breast, colon, prostrate, ovary, liver and brain, thus advocating its role in tumor suppression (66). Targeted repression of Cdh1 by E7 may, thus, be speculated as part of a mechanism designed to initiate neoplastic phenotype linked to HPV-associated cervical carcinogenesis.

Moreover, E7 considerably altered the expression of crucial Cdh1 substrates (Fig. 2D), alluding to the possible functions of such factors in E7-linked tumor maintenance. Potentially complementing our observations, past reports assert that loss of well-timed degradation of these molecules by Cdh1 is strongly connected to cell cycle abnormalities, genomic injury and poor prognosis in multiple tumors. Remarkably, we also discerned a rescue in FoxM1 from degradation imparted by Cdh1 following the introduction of E7. Embryogenic lethality in *foxm1* knockout mice model clearly demonstrates its importance in the survival of the host (67). In this aspect, up-regulation of this oncogenic transcription factor, as is common in a myriad of malignancies, is directly correlated with grade of the tumor and patient survival (68). FoxM1 knockdown, on the other hand, has shown promising results in effectively suppressing cancer development and progression (69, 70). Cdh1 is a master regulator of FoxM1, which ensures appropriate degradation of FoxM1 through recruitment of the APC/C E3 ubiquitin ligase (47). Although this mode of regulation has been extensively studied, the prospective impact of E7 on the Cdh1-FoxM1 relationship remains elusive. Our clear demonstration that E7 alleviates Cdh1 expression, culminating in elevated levels of FoxM1 (Fig. 6A, B), re-emphasizes that adequate regulation of FoxM1, *via* Cdh1, restricts eccentric oncogenic functions by tempering precise expression of FoxM1 target genes like Plk1, CyclinB1, Skp2, Aurora B kinase and CENP-A at the appropriate phases of cell cycle to facilitate an error-free cell cycle transition. When Cdh1 is abnormally degraded or rendered dysfunctional, as is the case in an E7 positive niche, this controlled governance is lost. The consequent atypical up-regulation of FoxM1 and its target molecules (Fig. 6A) can, therefore, be correlated as the contributing factors for inducing significant neoplastic features, as has been previously observed in cervical cancer cells (71). Essentially, we also noticed that the E7 mutants with defects in Cdh1 binding and subsequent destabilization failed to restore FoxM1 levels (Fig. 6D, E). Assessing the presence of such mutants in clinical specimens may shed more light on predicting the outcome of anti-cancer therapy targeted at E7-induced Cdh1 dysfunctioning. Also, our data suggest that E7 does not abolish the interaction between FoxM1 and Cdh1 but E7 binding to Cdh1 renders the latter inactive as well as incapable of degrading FoxM1 in spite of co-existence in the same trimeric complex (Fig. 6C). Although E7 was found to promote ubiquitin-induced degradation of Cdh1 (Figure 3F), detectable amount of Cdh1 was seen in association with FoxM1 in such E7-expressing cells. However, since we observed FoxM1 up-regulation in presence of E7 despite a positive FoxM1-Cdh1 complex formation, this raised several other potential mechanisms underlying E7-induced FoxM1 overexpression. These include sequestration of functional Cdh1 in the presence of E7, blocking the active site of Cdh1, thereby bequeathing FoxM1 inaccessible for degradation and formation of non-functional entity may also exist, which need to be addressed for future research. A noteworthy observation from previous independent reports is the loss of a tightly-regulated cell cycle as an aftermath of augmented accumulation of Plk1, Skp2 and others (72, 73). Our study demonstrated a novel mechanism depicting how E7 orchestrates an abnormal cell cycle transition through escalating the levels of Plk1, Skp2 and other downstream targets of FoxM1 by disrupting the functional dynamics of Cdh1 with its critical targets. Additionally, E7-mediated Cdh1 inactivation creates a cellular milieu conducive for viral DNA replication and pathogenesis through accumulation of cellular factors, Geminin, Cdt1 and FoxM1 that have implications in cancer development (Fig. 8).

It is of utmost importance to this study to peruse the biological fate of reduced levels of Cdh1 in establishment and sustenance of HPV-linked cervical tumor. Existing literature points to the ability of E7 to hijack a range of cell cycle checkpoints to favor an accelerated entry into S phase (74). Moreover, past studies indicate oncogenic features like unusual abundance of mitotic cyclins in the G1 phase and early onset of a prolonged S phase, leading to loss of cell cycle arrest despite assault to DNA replication fidelity, following Cdh1 inactivation (51, 75, 76). Our data was in agreement with such existing perceptions and further accomplished the deregulation of cell cycle by a dysfunctional Cdh1 as an effect of its transfiguration by E7. Cyclin-E is well-known for its imperative role in S phase entry and it is directly activated by HPV16 E7 (77). In addition, S phase cells with damaged DNA, in general, thwart the activation of CyclinB1, which is the prime regulator of M phase entry, to halt cells in G2 and inhibit neoplastic transformation. Contrary to this, HPV16 positive cells, despite having their DNA damaged, ensure over-expression of CyclinB1 for successful oncogenesis (78). Our results showing failure to mount an accumulation of CyclinB1 and CyclinE in presence of E7 with inability to inactivate Cdh1 (Fig. 7A, B) proposes the importance of Cdh1 in E7-related disruptions to the cell cycle. This postulate was further advocated by our BrdU assay and cell cycle analysis (Fig. 7C-F). E7’s inability to tweak Cdh1 activity caused a slow mitotic entry, which may result in corresponding reduction in cell proliferation and, thereby, tumorigenesis. Additionally, EMT has been implicated in aggressiveness of cancers and plays crucial roles in the tumor invasion process. Our finding that Cdh1-deregulation deficient mutant HPV16 E7 (1-15aa) or Cdh1 depletion in E7-expressing cells appreciably mitigated cervical cancer cell proliferation and metastasis accentuated the pivotal outcome of Cdh1 dysfunction in expediting E7-linked tumor progression (Fig. 8). Our depicted model supports this established paradigm wherein E7 reprograms the substrate specificities of cellular Cdh1, causing uncharacteristic cyclin expression, deviant cell cycle progression in addition to stimulated migratory and invasive potential of cervical carcinoma cells, which acts as a pre-requisite for E7-associated cellular transformation (Fig. 9).

**Fig. 9.**
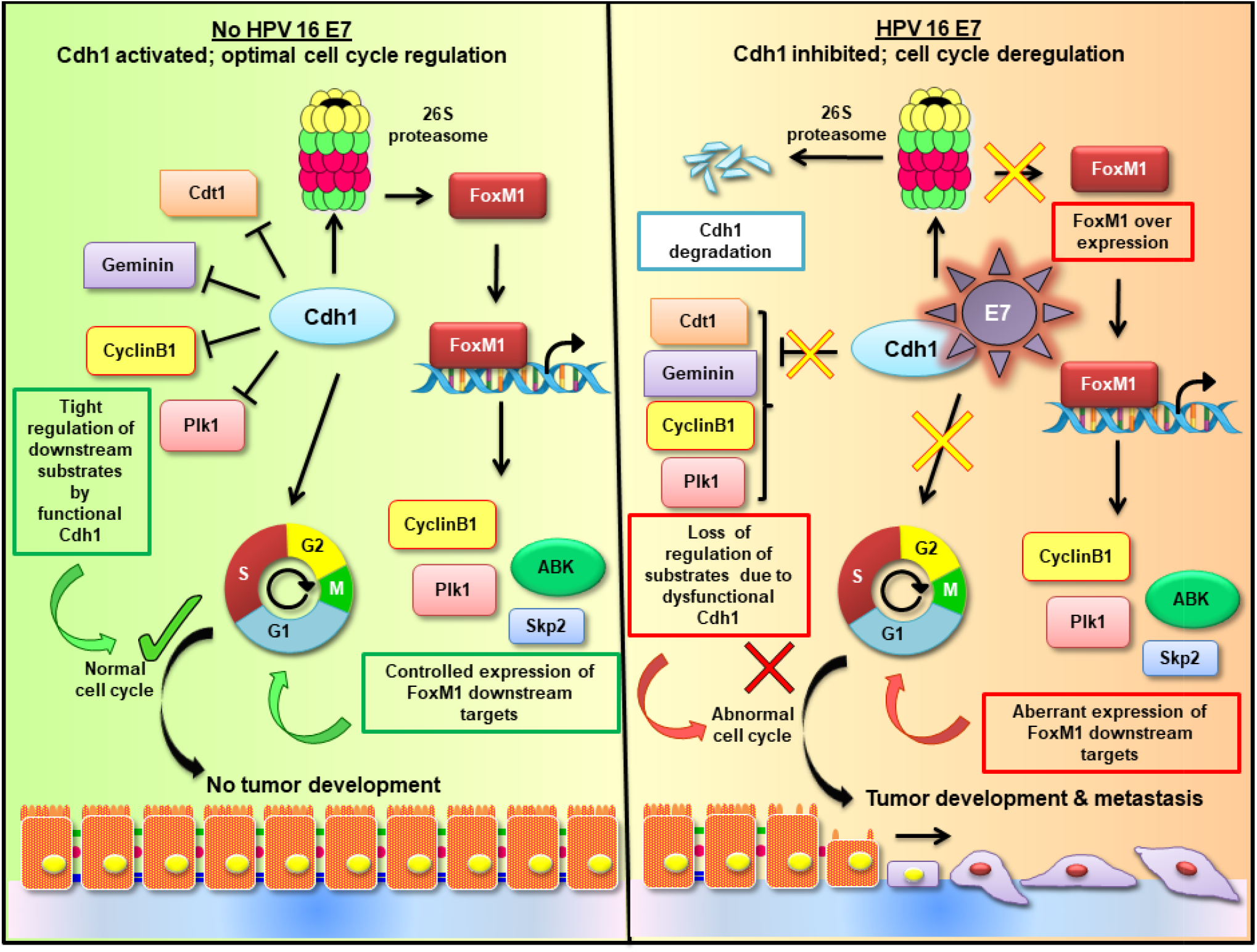
HPV16 E7 exploits the cellular APC-C/Cdh1 machinery to induce tumorigenic characteristics. Under typical conditions, Cdh1 maintains optimal levels of Geminin, Cdt1, CyclinB1 and Plk1 while targeting the oncogenic transcription factor FoxM1 for timely degradation by the 26S proteasome. This imparts a tight regulation on the transactivation of genes downstream to FoxM1, many of which are critical cell cycle regulators, CyclinB1, Plk1, Aurora B Kinase and Skp2. This cumulatively gives rise to the standard cell cycle transition. However, in cells infected with HPV, the E7 oncoprotein takes over this pivotal regulatory unit. Consequently, Cdh1 deregulation triggers irregular expression of all these genes, thus facilitating peculiar cell cycle progression in addition to other metastatic features such as EMT. Thus, E7-mediated modulation of Cdh1 poses as a pivotal point during HPV-related cellular transformation.

In summary, we have ascertained Cdh1 as a novel associating partner of E7 and reported new evidence that the destabilization of Cdh1 instilled by E7 undermined the controlled expression of critical downstream targets of Cdh1. This snowballed into early behaviors of neoplastic transformation, such as aberrant cell cycle transition, hyper-proliferation and metastasis. E7 mutants that are defective for Cdh1 interaction are impaired for these cancerous abilities. Thus, it is reasonable to assume that inhibiting the sabotaging of Cdh1 by HPV16 E7 may have therapeutic potential in HPV-induced carcinomas.

## EXPERIMENTAL PROCEDURES

### Antibodies and plasmid constructs

Monoclonal antibodies against the Flag tag epitope and Cdh1 were purchased from Sigma (St. Louis, MO) while antibodies against GFP, DsRED, Plk1, FoxM1, Rb, CyclinB1, CyclinE, β-Actin, GAPDH and HPV16 E7 were obtained from Santa Cruz Biotechnology Inc. (Santa Cruz, CA). Anti-Myc, anti-HA, anti-Skp2 and anti-Aurora B Kinase antibodies were purchased from Invitrogen (Carlsbad, CA). Antibodies against Cdt1, Geminin, CENP-A and Aurora A were obtained from Cell Signaling (Danvers, MA). Plasmid expressing Myc-Cdh1 was generously gifted by Pradip Raychaudhuri (UIC, USA). Plasmids expressing full length, wild type Myc-E7 and Flag-FoxM1 were constructed by in-frame cloning of PCR-amplified Myc-E7 and Flag-FoxM1, respectively, into mammalian expression vector pcDNA3.1, using Hind III and XhoI restriction sites. The single site mutations for replacing the cysteine residues at 24^th^ or 58^th^ position of E7 with glycine (p24Gly and C58Gly, respectively) were carried out *via* site-directed mutagenesis in Myc-E7 construct using the following pairs of primers: 5’-GAG ACA ACT GAT CTC TAC GGT TAT GAG CAA TTA AAT GAC-3’ and 5’-GTC ATT TAA TTG CTC ATA ACC GTA GAG ATC AGT TGT CTC-3’, 5’-TAC AAT ATT GTA ACC TTT GGT TGC AAG TGT GAC TCT ACG-3’ and 5’-CGT AGA GTC ACA CTT GCA ACC AAA GGT TAC AAT ATT GTA-3’. DsRED-E7 construct was prepared by sub- cloning full length E7 into pDsRED-Express-N1, utilizing Hind III and Apa1 restriction sites. Plasmids encoding Flag-Cdh1 and EGFP-Cdh1 were generated by in-frame cloning of PCR amplified Flag-Cdh1 and Cdh1 into Hind III and EcoR1 sites of pcDNA3.1 and pEGFP-C1 vectors (Clonetech laboratories, Inc.), respectively. Plasmids expressing DsRED-E7 (wild type) and its deletion mutants (1-15, 16-37 and 38-98aa), EGFP-Cdh1, Flag-Cdh1 (wild type) and its deletion mutants (1-124, 1-200 and 201-493aa) were cloned using the following primers:

DsRED-E7: 5’-GAGAAGCTTATGCATGGAGATACACCT - 3’ and 5’- GAGGGGCCCTTGGTTTCTGAGAACAGAT - 3’;

DsRED-E7 (1-15aa): 5’- GAGAAGCTTATGCATGGAGATACACCT - 3’ and 5’-GAGGGGCCCACAAATCTAACATATATT - 3’;

DsRED-E7 (16-37aa): 5’ - GAGAAGCTTATGCAACCAGAGACAACTGATCTC - 3’ and 5’ - TCAGGGCCCATTCATCCTCCTCCTC - 3’;

DsRED-E7 (38-98aa): 5’ - GAGAAGCTTATGATAGATGGTCCAGCT - 3 and 5’-GAGGGGCCCTTGGTTTCTGAGAACAGAT - 3’;

EGFP-Cdh1: 5’- GAGAAGCTTATGACCAGGACTATGAGCGG - 3’ and 5’ -GAGGAATTCTTACCGGATCCTGGTGAA - 3’;

Flag-Cdh1 (wt): 5’- GGAAAGCTTGACCAGGACTATGAGCGG - 3’ and 5’- GGAGAATTCTTACCGGATCCTGGTGAA - 3’;

Flag-Cdh1 (1-124aa): 5’- GGAAAGCTTGACCAGGACTATGAGCGG - 3’ and 5’-GGAGAATTCTTACTTCTCAGGCGTGGAGGG - 3’;

Flag-Cdh1 (1-200aa): 5’- GGAAAGCTTGACCAGGACTATGAGCGG - 3’ and 5’-GGAGAATTCTTACACATTGAGGGACGACCA - 3’;

Flag-Cdh1 (201-493aa): 5’- GGAAAGCTTCTCAGCGTGGGGCTAGGC - 3’ and 5’-GGAGAATTCTTACCGGATCCTGGTGAA - 3’.

Sources of plasmids expressing HA-Ub, EGFP-FoxM1 and pSG5-E7 have been described in our earlier publications (27).

### Cell culture, Transfection and Western blotting

HEK293T, Phoenix Ampho and C33A cells were cultured in DMEM medium (Invitrogen, Carlsbad, CA) and CaSki cells were grown in RPMI medium (Invitrogen, Carlsbad, CA), both supplemented with 10% fetal bovine serum (Invitrogen, Carlsbad, CA), 100 U/ml of penicillin and 100 mg/ml streptomycin (Invitrogen, Carlsbad, CA) in a humidified incubator with 5% CO_2_ atmosphere at 37°C, as described previously (27). For over-expression experiments, HEK293T cells were transiently transfected by calcium phosphate method as described previously (79). C33A cells were transfected with either Lipofectamine 2000 (Invitrogen, Carlsbad, CA) or FuGENE transfection reagent (Roche Diagnostics, Indianapolis, IN) as per the manufacturers’ instructions. The medium was changed 16 h post transfection and 24 h thereafter, cells were lysed in lysis buffer containing 50 mM Tris-HCl, 400 mM NaCl, 0.2% Nonidet P-40, 10% glycerol and protease inhibitor cocktail (Roche Applied Science, Mannheim, Germany). Equal amounts of whole cell lysates were processed for SDS-PAGE and subjected to immunoblotting with appropriate antibodies.

### siRNA transfection

C33A-E7 stable cells were transfected with siRNA specific for Cdh1, purchased from Sigma (St. Louis, MO), at a final concentration of 100 nM using Oligofectamine (Invitrogen, Carlsbad, CA). A scrambled siRNA was used as a negative control.

### Generation of stable cell line

C33A with stable expression of HPV16 E7 were generated by infecting HPV negative C33A cell line with pBabe-E7-puromycin based retroviral vector and selected with puromycin. Recombinant retroviruses were produced in Phoenix Ampho packaging cell line. Infection of 50% confluent C33A was performed with viral supernatant for 4-6 times. Pure, virally transduced population was selected and maintained in media containing puromycin (0.5 μg/ml). Over-expression was verified by assessing E7 at RNA level by RT-PCR. For generation of HPV16 E7 knockdown cells, CaSki cells were transduced with shE7 (sense: 5’-GACAGAGCCCAUUACAAUA - 3’ and anti-sense: 5’-UAUUGUAAUGGGCUCUGUC-3’) retroviral supernatants. Knockdown was verified by assessing the expression of endogenous E7 using Western analysis.

### RT-PCR

Total RNA was isolated by using Trizol reagent (Sigma, St. Louis, MO), according to the manufacturer’s instructions. PCR was performed using one step SuperScript^R^ RT-PCR kit (Invitrogen, Carlsbad, CA) with E7 specific primers: 5’-GAGAAGCTTCATGGAGATACACCTACATTG-3’ (sense primer) and 5’-GCGCTCGAGTTATGGTTTCTGAGAACAGAT-3’ (anti-sense primer) and GAPDH primers 5’-ACCTGACCTGCCGTCTAGAA-3’ (sense primer) and 5’-TCCAACCACCCTGTTGCTGTA-3’ (anti-sense primer) as per the manufacturer’s protocol.

### Cycloheximide chase assay

Protein degradation assay was performed as described previously (80). In brief, cells were transfected with the desired expression vectors and 16 h post transfection, they were trypsinized, pooled and reseeded equally in petri plates. After growing for another 16 h, cells were treated with 100μg/ml cycloheximide (CHX: Sigma, St. Louis, MO), harvested at varied time intervals and equal amounts of the whole cell lysates were subjected to Western blot analysis. Densitometric analyses of scanned images were carried out using Multi Gauge V3.0 (Fujifilm, Tokyo, Japan) software.

### Co-immunoprecipitation

Cell extracts were prepared in lysis buffer containing 150 mM NaCl, 1% NP-40, 5 mM EDTA, 50 mM Tris-HCl (pH 7.5), 2 mM PMSF (phenylmethanesulfonyl fluoride), 2 mM NaF, 1 mM Na_3_VO_4_ and protease inhibitor cocktail, as described in our previous publication (27). Briefly, lysates were pre-cleared with 10 μl protein-G bead slurry (50%) at 4°C with constant rocking for 1 h. For IP, 1.5 mg pre-cleared extract was gently rocked with 1 μg of antibody for 4 h at 4°C. Thereafter, 40 μl of protein-G bead suspension was added to the above mixture and allowed to rock for an additional 1 h at 4°C. The beads were washed 3-4 times with the lysis buffer and the bound protein was eluted with SDS-PAGE sample dye. Beads were spun down and the clear supernatant was resolved by SDS-PAGE and subjected to immunodetection with appropriate antibodies.

### *In vivo* ubiquitination assay

HEK293T cells were transfected with vectors expressing Flag-Cdh1, HA-Ub and Myc-E7 or other constructs as indicated in the figures, using calcium phosphate method. As discussed in our earlier publication (80), after 36 h of transfection, cells were treated with 20 μM MG132 (Sigma, St. Louis, MO) for 5 h. MG132 (Z-Leu-Leu-Leu-al) is a peptide aldehyde, which effectively blocks the proteolytic activity of the 26S proteasome complex (81). It is a specific, potent, reversible, and cell-permeable proteasome inhibitor. Following transfection, cells were lysed in lysis buffer containing 150 mM NaCl, 1% NP-40, 1 mM EDTA, 40 mM HEPES (pH 7.0), protease inhibitor cocktail, 2 mM PMSF, 2 mM NaF, and 1 mM Na_3_VO_4_. Lysates were pre-cleared with 10 μl protein G-beads for 1 h at 4°C. Thereafter, IP against the Flag epitope was performed by incubating 2 mg of pre-cleared supernatant with 1 μg antibody for overnight at 4°C. The unbound proteins were removed by washing the beads extensively with modified RIPA buffer (82) containing 150 mM NaCl, 1% NP-40, 1% deoxycholate, 0.05% SDS, 1 mM EDTA, 40 mM HEPES (pH 7.0) and 2 mM PMSF. Protein complexes bound to beads were resolved by SDS-PAGE and detection was carried out using appropriate antibodies.

### FACS analysis

Cell cycle analysis was conducted as described before (80). Briefly, logarithmically growing C33A cells were seeded, followed by either transfection or knockdown as indicated. Cells were grown for 48 h in a humidified CO_2_ incubator at 37°C. Cells were harvested using PBS containing 4 mM EDTA, washed with PBS and fixed in ice cold 85% ethanol for overnight at −20°C. Cells were thereafter washed in PBS and re-suspended in 0.5 ml of PBS containing 1 mg/ml RNaseA (Himedia, India) and 150 μg/ml propidium iodide (Sigma, St. Louis, MO). After incubation at RT for 30 mins in dark conditions, cells were acquired on a flow cytometer using FACSCalibur™ (BD Biosciences, San Jose, CA). At least 20,000 cells were counted for each sample. Cell cycle profile was determined by using the CellQuest cell analysis software package.

### Fluorescence microscopy

HEK293T cells were cultured in DMEM along with coverslips and transiently transfected with plasmids expressing either DsRED-E7 or EGFP-Cdh1 alone and together. Forty hours after transfection, coverslips were fixed in 4% *para*-formaldehyde solution (w/v) in phosphate buffer saline for 20 mins at RT, followed by additional PBS washes. Permeabilization was carried out in 0.5% triton in PBS for 20 mins at RT, after which cells were washed and counterstained with DAPI (4’,6-diamidino-2-phenylindole). Coverslips were mounted on microscopic slides and images were acquired using fluorescence microscope (Nikon-Eclipse-Ti-S, Tokyo, Japan).

### Confocal microscopy

Cells transfected with either EGFP-Cdh1 alone or in combination with DsRED-E7 WT and mutants were fixed in 4% *para*-formaldehyde for 20 mins at RT, washed thrice with PBS and stained with 1 μg/ml DAPI for 5 mins at RT. Cells were washed again and mounted in ProLong Gold Anti-fade reagent (Molecular Probes, Eugene, OR) and captured using Leica SP-5 confocal laser-scanning microscope in CIF, UDSC.

### BrdU Immunofluorescence

BrdU assay was carried out using the protocol published earlier (80). Briefly, cells were seeded with coverslips in a 12-well plate so that cells attain around 80% confluency the following day and allowed to grow in a 37°C humidified CO_2_ incubator overnight. The following day, BrdU solution was added to the cells and allowed to incubate for 1 h at 37°C. Cells were washed twice with 1X PBS followed by fixation with 4% *para*-formaldehyde. Cells were washed thrice with 1X PBS and incubated in 1X PBS-T (1X PBS containing 0.1% Triton X 100) for permeabilization. Following washes, cells were incubated with 1(N) HCl on ice for 10 mins followed by treatment with 2(N) HCl at RT for 10 mins. Phosphate citric acid buffer (pH 7.4) was added and cells were incubated at RT for 10 mins followed by washes with 1X PBS-T thrice. Cells were then blocked in 5% BSA for 1 h at 4°C. Cells were then incubated overnight with anti-BrdU antibody (RPN202, GE Healthcare, Buckinghamshire, UK) at 4°C. Following day, cells were washed thrice with 1X PBS-T and incubated with anti-mouse Alexa Fluor 594 conjugated secondary antibody (Invitrogen, Carlsbad, CA) for 1 h at RT. Cells were given three washes with 1X PBS and nuclei were counter stained with DAPI. Cells were mounted on slides and observed under fluorescence microscope (Nikon-Eclipse-Ti-S, Tokyo, Japan). The number of BrdU positive cells was counted.

### Migration and invasion assays

Cells were seeded to complete confluence in a monolayer in a 24-well plate for wound healing assay. A wound was created by scratching firmly with a 20 μl tip and baseline (time zero) images were captured. Cell were incubated at 37°C in a humidified incubator with 5% CO_2_ and photographed at the indicated time-points and the percent wound remaining was measured and plotted. Invasion assay was performed using 24-well BD BioCoat Matrigel invasion chambers (BD Biosciences, San Jose, CA) according to the manufacturer’s instructions. Briefly, inserts were first rehydrated using DMEM containing 10% FBS for a final volume of 0.5 ml in the top inserts and 0.75 ml in the wells of a 24-well dish (Corning Inc., Corning, NY) for at least 2 h. Following this, the medium was removed from the upper inserts and a total of 1×10^5^ cells were plated in 0.5 ml of serum free medium. Cells were allowed to invade through the matrigel towards the FBS containing medium, which acts as a chemoattractant, for 48 h. Non-invading cells were removed using a damp cotton swab while the invading cells were fixed in methanol for 10 mins and stained with 0.5% crystal violet (Sigma, St. Louis, MO) stain. Cells in at least six randomly selected fields were then counted under a light microscope.

### Statistical analysis

Results were expressed as means ± standard errors of the means (SEM). Statistical significance was assessed by analysis of variance and was defined as *, *P* < 0.05 and *P* > 0.01; **, *P* < 0.01 and *P* > 0.001; ***, *P* < 0.0001.

## ACKNOWLEDGEMENTS

This work was supported by research grant to AN funded by Council of Scientific and Industrial Research (CSIR no. 37 (1682)/17/EMR-II), Department of Biotechnology (DBT no. BT/PR15422/MED/30/1705/2016), University Grants Commission (UGC-SAP Program), Department of Science and Technology (DST)-PURSE (RC/2014/7114) and Institute of Eminence of Delhi University. The funders had no role in the experimental designs or data collection and analysis or the decision to submit the work for publication. The DBT is thanked for providing fellowship to NJ and DN while CSIR is acknowledged for granting fellowship to PSC. Central Instrumentation Facility, UDSC is duly acknowledged for providing common infrastructure.

