## Supplementary files for "The Anaphase Promoting Complex/ cyclosome co-activator, Cdh1, is a novel target of human Papillomavirus 16 E7 oncoprotein in cervical oncogenesis"

**Supplementary Information**

**Supplementary Table 1.**

|  | **Colocalization** | | | |
| --- | --- | --- | --- | --- |
|  | **E7 WT** | **E7 1-15aa** | **E7 16-37aa** | **E7 38-98aa** |
| **Pearson’s correlation** | 0.5435 | 0.3539 | 0.6638 | 0.6061 |
| **Overlap co-efficient** | 0.5051 | 0.5067 | 0.7001 | 0.7057 |
| **Colocalization rate** | 68.3% | 39.42% | 43.39% | 79.96% |

**Supplementary Fig. 1** Representative scatter plot depicting the co-localization between EGFP-Cdh1 and DsRED-E7 WT or mutants 1-15aa, 16-37aa and 38-98aa.

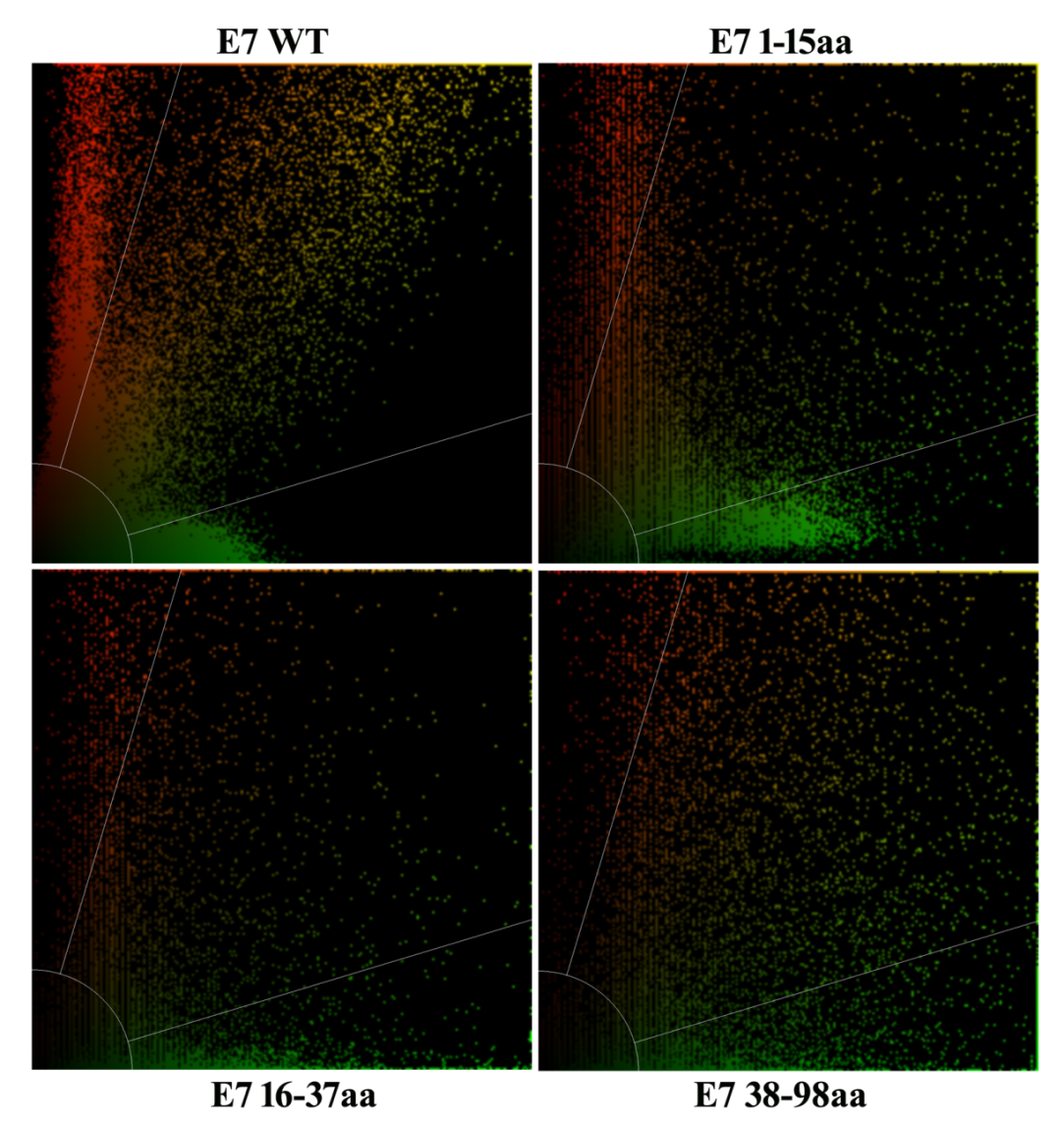
